# *Sod1* trisomy causes ENS developmental defects and susceptibility to Hirschsprung disease via neuronal *Ret* suppression and glial remodeling

**DOI:** 10.64898/2026.01.26.701837

**Authors:** Gabriel Grullon, Jarod Rollins, Lauren Wilkes, Aamir Zuberi, Aravinda Chakravarti, Sumantra Chatterjee

## Abstract

Down syndrome (DS; trisomy 21) confers a ∼100-fold increased risk of Hirschsprung disease (HSCR), yet the causal contributions of specific chromosome 21 genes remain unresolved. Here we show that increased dosage of SOD1 alone is sufficient to perturb enteric nervous system (ENS) development. We engineered a humanized *SOD1* trisomic mouse line by inserting a 28 kb human *SOD1* locus into ROSA26 and genetically profiled the distal colon at postnatal day 0 using single-cell RNA-seq and immunofluorescence. *Sod1/SOD1* was elevated ∼1.5X in the ENS but varied by cell type, with transcriptionally active progenitors showing the greatest increase. Cell composition shifted toward transcriptionally active cells and glia, with concomitant loss of excitatory and inhibitory motor neurons and interneurons. Genetically, *Sod1/SOD1* trisomy downregulated synaptic and neuronal communication programs but upregulated DNA replication/cell-cycle and genome maintenance pathways, especially within glia. Consequently, key HSCR genes were dysregulated: *Ret*, *Ednrb*, and *Sema3c* were decreased, while *Sema3a* ,a negative guidance cue, was increased. *Ret* was selectively reduced in inhibitory and excitatory motor neurons and progenitors, unchanged in glia, and reduced at the protein level *in vivo*. Within glia, Sod1/SOD1 was particularly elevated in proliferating/active glia with a glia-specific bias toward endogenous mouse *Sod1* expression. Taken together, these data support a dual mechanism whereby increased *Sod1/SOD1* dosage suppresses *RET*-dependent neurogenesis while independently promoting reactive/proliferative glial states. Thus, *SOD1* is sufficient to alter ENS development significantly and provide the susceptibility substrate for HSCR with further reductions in *RET* gene expression leading to aganglionosis.

## Introduction

Down syndrome (DS), the commonest genetic cause of intellectual disability, results from trisomy of human chromosome 21 (HSA21) with incidence increasing with maternal age (Antonarakis et al., 2020; Hook et al., 1983). T21 affects multiple organ systems, and individuals with DS are predisposed to over 40 congenital anomalies—including cardiovascular, gastrointestinal, musculoskeletal, immune, and neurological defects—yet paradoxically are protected from most solid tumors.(Torfs and Christianson, 1998) Despite extensive investigation the underlying pathophysiology of DS-associated traits remains poorly understood. These no doubt arise from increased dosage of *specific but largely unknown* gene(s) on chromosome 21, however, whether the source of phenotypic variation within DS is from *sequence variation* in HSA21 *primary* genes or non-HSA21 *modifier* genes and/or *interaction* between HSA21 trisomy and non-HSA21 genetic variation is unknown. Preliminary evidence suggests that all of these mechanisms can exist (Arron et al., 2006; Bushman et al., 2015; Ono et al., 2015; Wechsler et al., 2002; Wiseman et al., 2018).

One of the most striking DS-associated phenotypes is Hirschsprung disease (HSCR), or congenital intestinal aganglionosis, which occurs at a 100-fold elevated incidence of 13.8 per 1,000 live births in DS as compared to 14 per 100,000 in the general population (Amiel et al., 2008; Badner et al., 1990). However, T21 alone is not sufficient to cause HSCR, indicating that additional genetic or epigenetic modifiers, whether on chromosome 21 or elsewhere, must interact with the trisomic background to confer disease risk (Arnold et al., 2009).

Although more than two dozen genes predispose to HSCR, its major genes encode the receptor tyrosine kinase *RET* and the G-protein coupled receptor *EDNRB(*Tilghman et al., 2019). Disease-associated mutations lead to reduced expression of *RET* and *EDNRB* which are rate-limiting genes in a gene regulatory network (GRN) controlling survival, proliferation, and migration of enteric neural crest-derived cells (ENCDCs) and enteric neurogenesis (Chatterjee and Chakravarti, 2019; Chatterjee et al., 2016; Fries et al., 2025). Thus, it was intriguing that in 2007 de Pontual and colleagues identified a genetic association between *RET* and a number of HSCR-associated syndromes, including DS(de Pontual et al., 2007), a finding confirmed by DS and control family analyses by Arnold and colleagues (Arnold et al., 2009). The specific common variant association was later recognized to be within a gut-specific transcriptional enhancer of *RET* that *reduced* its gene expression leading to increased HSCR genetic susceptibility(Chatterjee et al., 2023; Chatterjee et al., 2016). These findings taken together lead to our hypothesis that increased dosage of one or more HSA21 gene interacts with *RET* to further reduce its gene expression in individuals who already harbor the hypomorphic *RET* enhancer variant leading to <50% *RET* gene expression and aganglionosis(Fries et al., 2025). Note, Other gastrointestinal malformations such as esophageal and duodenal atresia, imperforate anus, and gut malrotation are also overrepresented in DS, suggesting shared developmental origins (Torfs and Christianson, 1998).

In this study, we demonstrate that *SOD1* on HSA21 is one such gene. Our prior studies of both mouse models of HSCR, harboring coding and enhancer loss-of-function mutations in *Ret* (Chatterjee et al., 2019; Fries et al., 2025; Vincent et al., 2023), and in human embryonic gastrointestinal tracts deficient in *RET* (Chatterjee et al., 2023), have identified eight HSA21 genes that are significantly dysregulated in both species: *S100B, SOD1, ADAMTS1, PRMT2, CBR1, PDXK, PCP4*, and *NDUFV3,* all of which are upregulated except *ADAMTS1*. Among these, *SOD1* stands out as a particularly compelling HSCR-interacting candidate gene. First, mutations and overexpression of *SOD1* are causally linked to familial forms of amyotrophic lateral sclerosis (ALS), where toxic gain-of-function effects lead to motor neuron degeneration through oxidative stress, protein aggregation, and disruption of intracellular signaling pathways (Rosen et al., 1993; Tu et al., 1996). Second, beyond its canonical role as a cytoplasmic antioxidant enzyme, *SOD1* has also been shown to interfere with receptor-mediated signaling pathways: notably, overexpression of SOD1 impairs RET dimerization (Kato et al., 2000). Here we utilize mouse models to demonstrate that *SOD1* overexpression can diminish *RET* function, creating a permissive environment for ENS developmental failure even in the absence of pathogenic *RET* variants.

To be sure, other HSA21 genes may also contribute to the elevated HSCR risk in DS. Previous studies of individuals with segmental or partial trisomy 21 had suggested that a ∼13 Mb critical region of HSA21 was potentially involved in HSCR susceptibility (Korbel et al., 2009), with *DSCAM* as a candidate gene based on HSCR association studies (Jannot et al., 2013). To gather corroborating evidence, others have studied the DS mouse models Ts65Dn and Tc1 that are trisomic for many genes orthologous to HSA21 including *Dscam* and *Dyrk1a*, however, neither model demonstrate any defects in enteric neurogenesis at embryonic day (E)12.5 and retain a largely intact myenteric plexus architecture in adulthood (Schill et al., 2019). Interestingly, Ts65Dn mice display a reduction in submucosal neurons, but this defect persists even when copy number for *Dscam* or *Dyrk1a* is normalized. These equivocal findings are in contrast to the *SOD1*-dependent defects in enteric neurogenesis we demonstrate.

Progress in understanding DS-associated phenotypes require the study of specific mouse models with perturbations of one gene at a time. To this end, we generated a humanized trisomic *SOD1* mouse model by inserting a 28 kb human genomic fragment encompassing the *SOD1* gene and its native regulatory elements into the mouse Rosa26 locus, ensuring stable, ectopic expression without disrupting endogenous gene networks. In this model, we demonstrate a selective downregulation of *Ret* in enteric neurons at postnatal day 0 (P0) with the most pronounced reduction in inhibitory motor neurons, the neuronal subtype deficient HSCR pathogenesis (Fries et al., 2025; Vincent et al., 2023), accompanied by a decrease in the number of mature neurons within the ENS. In contrast, we observed a marked expansion of glial subpopulations characterized by proliferative or reactive states, accompanied by a reduction in differentiated glial types such as astrocytes and microglia. These findings demonstrate that *SOD1* overexpression alone may be sufficient to drive cell-type–specific disruptions in the developing ENS. While neuronal defects appear to arise via *RET* suppression, glial changes reflect a distinct pathway of dysregulation. This dual mechanism provides a compelling explanation for how trisomy 21 may predispose individuals with Down syndrome to HSCR, through both loss of neurogenic capacity and dysregulated gliogenesis.

## Results

### Sod1 trisomy induces gene expression changes in ENS-associated pathways in the distal colon

To model the effect of *SOD1* trisomy *in vivo*, we engineered a mouse line (*TS-SOD1*) carrying a 23.8 kb segment (chr21:31,653,123-31,676,942; GRCh38) of the human genome containing the *SOD1* gene and its native regulatory context, inserted into the mouse *Rosa26* locus (**Figure 1A**). This approach avoids disruption of endogenous loci while enabling stable expression of an additional *SOD1* allele.

**Figure 1:**
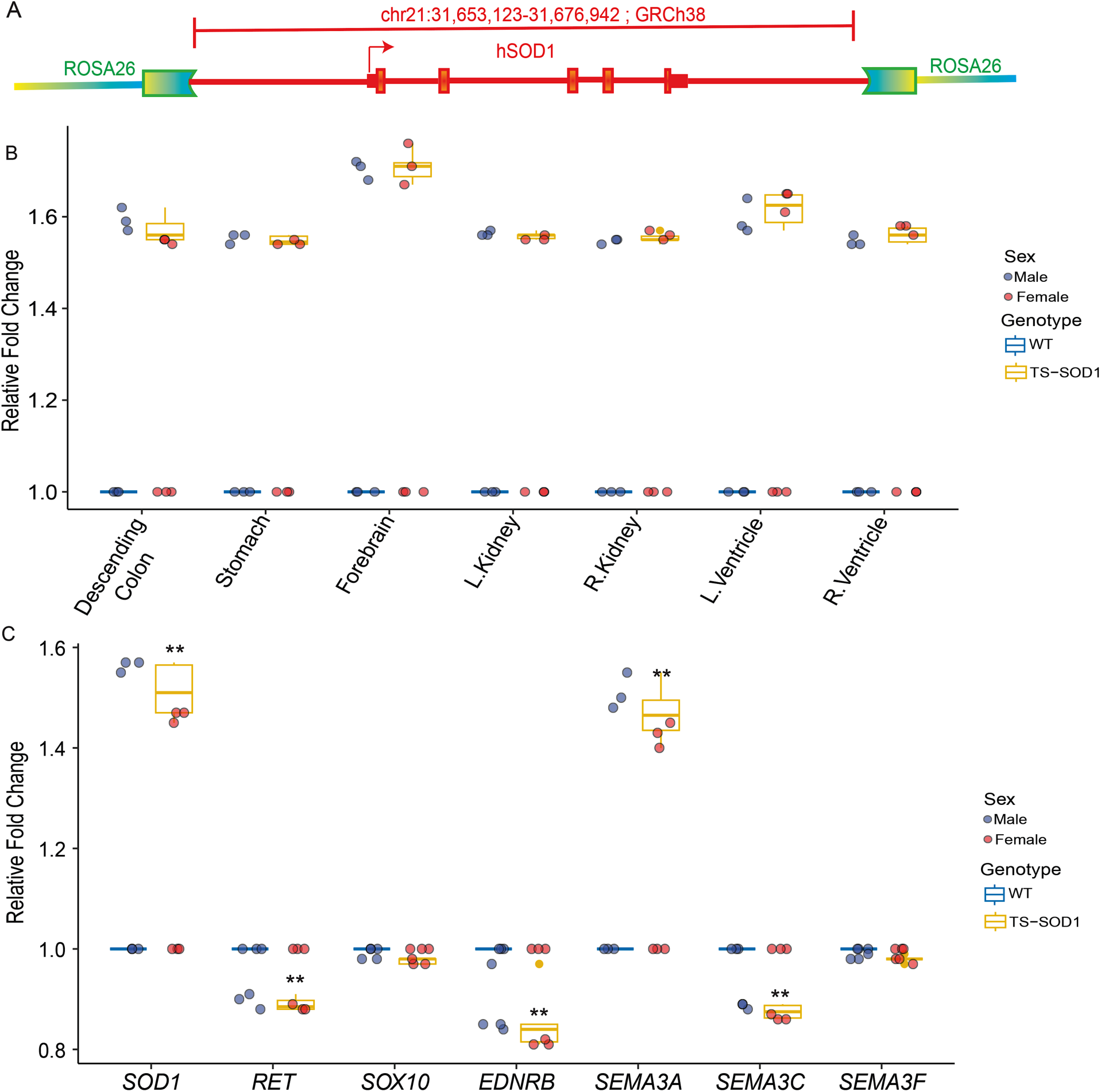
A humanized *Sod1/SOD1* trisomic mouse model shows elevated *Sod1/SOD1* gene expression and dysregulation of enteric nervous system genes associated with Hirschsprung disease. (A) A schematic of the ∼23.8 kb human genomic fragment containing *SOD1* and its regulatory elements (chr21:31,653,123-31,676,942; GRCh38) inserted into the mouse ROSA26 locus to generate the trisomic mouse line. (B) Quantitative RT-PCR demonstrating tissue-wide upregulation of *Sod1/SOD1* gene expression in TS-*Sod1/SOD1* neonates (P0) relative to wild-type (WT) littermates. Notably, expression in the forebrain reaches ∼1.7-fold, while distal colon, stomach, left and right kidneys and left and right ventricles of the heart show ∼1.5-fold elevation, consistent with trisomic dosage. (C) Gene expression analysis of the distal colon at P0 reveals significant dysregulation of Hirschsprung disease (HSCR)–associated genes in TS-*Sod1/SOD1* mice. Expression of *Ret*, *Ednrb*, and *Sema3c* are significantly reduced, while *Sema3a* is upregulated ∼1.5-fold relative to WT controls. The results in (B) and (C) are mean ± SEM with p < 0.01 by two-way ANOVA with genotype and sex as factors.

We confirmed that *TS-SOD1* neonates exhibit elevated *Sod1* expression across multiple organs at postnatal day 0 (P0) by quantitative RT-PCR (**Figure 1B**). All tissues, including the distal colon, kidneys, stomach, and ventricles of the heart, displayed 1.5-fold upregulation (p<0.001 for all) compared to wild-type littermates, consistent with gene dosage expectations. Notably, *Sod1* expression in the forebrain was further elevated, reaching a mean ∼1.7-fold increase (p<0.001), suggesting possible region-specific enhancer effects or differential chromatin accessibility at the *Rosa26* locus.

We next examined whether *Sod1* trisomy perturbs expression of genes from 3 signaling pathways critical for enteric nervous system (ENS) development and Hirschsprung disease (HSCR) susceptibility in the distal colon, namely, the receptor tyrosine kinase- *Ret*, the G protein coupled receptor- *Ednrb* and axon guidance molecules- Semaphorins (Fries and Chatterjee, 2024). Compared to WT littermates, *TS-SOD1* mice showed significant downregulation of *Ret (*0.5-fold, p<0.01), *Ednrb* (0.8-fold, p<0.01), and *Sema3c (*0.6-fold, p<0.01), all of which play essential roles in enteric neural crest cell migration, proliferation, or axon guidance (**Figure 1C**). In contrast, *Sema3a* expression was markedly increased (∼1.5-fold, p<0.01), indicating that *Sod1* trisomy may induce both suppressive and activating effects on key ENS regulatory pathways. It has been previously demonstrated that *Sema3a* functions as a negative regulator of enteric neural crest–derived cell (ENCDC) migration, as mice lacking *Sema3a* exhibit premature entry of sacral neural crest–derived cells into the hindgut (Anderson et al., 2007). Therefore, overexpression of this negative guidance cue may inhibit ENCDC migration, contributing to cellular loss in the developing enteric nervous system.

These findings demonstrate that Sod1 overexpression alone—among genes on chromosome 21—is sufficient to disrupt the transcriptional program of the developing gut, affecting key regulators of enteric nervous system development and Hirschsprung disease susceptibility.

### Sod1 trisomy perturbs ENS cell-type composition, differentiation trajectories, and cell-type-specific expression patterns in the neonatal distal colon

To investigate the impact of Sod1 trisomy on enteric nervous system (ENS) development, we performed single-cell RNA sequencing (scRNA-seq) on distal colonic tissues isolated from postnatal day 0 (P0) TS-SOD1 and wild-type (WT) littermates. The distal colon, which is most frequently affected in Hirschsprung disease, was dissected and dissociated for unbiased profiling of the ENS cellular landscape. Uniform Manifold Approximation and Projection (UMAP) of transcriptomes from 3879 (2288 from wild-type and 1591 from TS-SOD1 mice) individual cells revealed genotype-specific distributions, indicating underlying differences in cellular composition and/or gene expression (**Figure 2A**).

**Figure 2:**
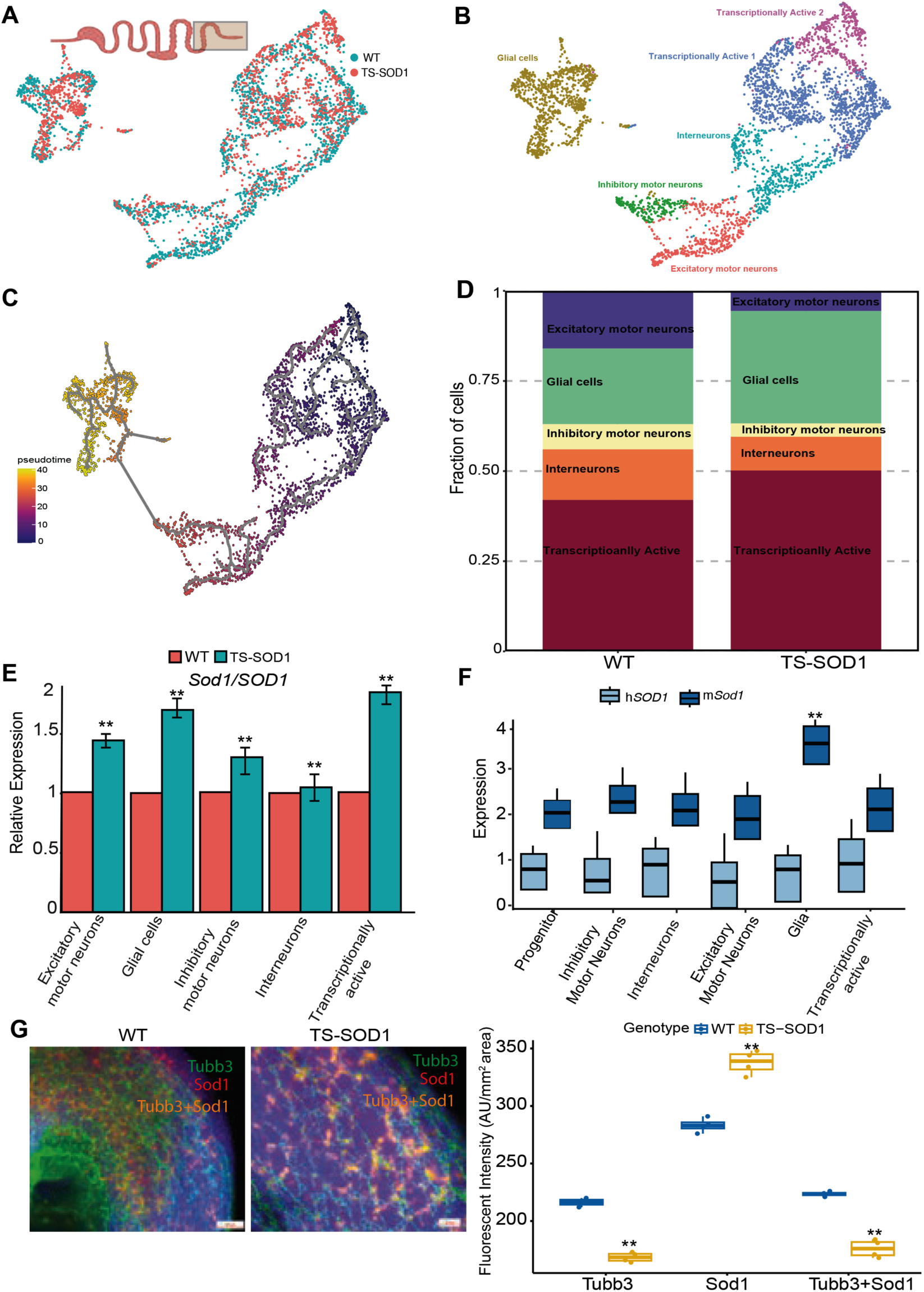
Single-cell transcriptome analysis reveals altered ENS cell composition and *Sod1/SOD1* expression dynamics in the TS- *Sod1/SOD1* neonatal distal colon. (A) UMAP of scRNA-seq data from the distal colon at P0 showing separation of WT and trisomic cells. (B) Cell-type annotation based on marker gene expression identifies five major ENS-derived populations: transcriptionally active progenitors, interneurons, inhibitory motor neurons, excitatory motor neurons, and glial cells. (C) Differentiation trajectory as inferred by *Monocle* (Trapnell et al. 2014) shows glial cells as fully differentiated and distinct from neuronal lineages. Excitatory and inhibitory motor neurons occupy parallel pseudo-temporal branches, reflecting coordinated maturation. (D) Plot of relative cell type abundance across genotypes. All mature neuronal types and interneurons are reduced in TS-*Sod1/SOD1* animals whereas transcriptionally active cells and glia are significantly increased. (E) Expression of *Sod1/SOD1* across ENS cell types. All show elevated and at least 1.5-fold increased expression in cells from trisomic than disomic animals except for interneurons (∼1.2x). (F) The ratio of endogenous mouse *Sod1* to human *SOD1* transcripts is generally 2:1 as expected but glial cells exhibit a significantly higher ratio of 3.2. (G) Immunofluorescence quantification in distal colon sections stained for *Tubb3* (neuronal marker) and *Sod1*. TS-*Sod1/SOD1* mice show ∼1.3-fold reduction in *Tubb3* signals and ∼1.2-fold increase in *Sod1*. Co-expressing cells (*Tubb3* and *Sod*1) exhibit ∼1.25-fold lower intensity in TS-*Sod1/SOD1* mice vs. WT. Data are shown as mean ± SEM; statistical significance (p < 0.01) was determined using two-way ANOVA or unpaired t-tests.

From these data we identified 1,423 high-variability genes and 15 principal components (PCs) that explain 90% variance, to cluster cells based on their differential gene expression (FDR<0.05) across clusters, relative to the mean expression in other clusters. We used marker genes to specify the identity of each cluster as follows: transcriptionally active progenitor-like cells (*Sox10, Ednrb*, *Hoxd5 and Hoxd6)* interneurons (*Scrt1, Nxph4 and Dlx5*), inhibitory motor neurons (*Nos1* and *Vip)*, excitatory motor neurons (*Calb2* and *Chat)*, and glial cells (*S100b, Gfap, Plp1*) (**Figure 2B**). These clusters broadly reflected known developmental intermediates and terminally differentiated subtypes of the ENS at this stage (Elmentaite et al., 2021; Fries et al., 2025; Majd et al., 2024; Vincent et al., 2023). The presence of transcriptionally active cells was indicative of ongoing neurogenesis, while the separation of neuronal and glial populations reflected the expected lineage bifurcation (Guyer et al., 2023; Laddach et al., 2023).

To better understand lineage dynamics, we performed trajectory analysis using Monocle (Trapnell et al., 2014). This revealed a clear pseudotemporal axis of differentiation, with glial cells (*Plp1, Mpz, S100b)* occupying a distinct terminal branch, suggesting complete divergence from the neuronal lineage by P0 (**Figure 2C**). In contrast, inhibitory and excitatory motor neurons emerged along parallel arms within a shared trajectory, suggesting simultaneous or convergent maturation paths. This organization reflects coordinated differentiation of neuronal subtypes from a common progenitor pool and provides a framework to assess how *Sod1* dosage affects these transitions.

Next, we quantified the relative abundance of each ENS cell type in TS-SOD1 and WT mice. All mature neuronal subtypes, including interneurons (<15%, p<0.001), inhibitory motor neurons (<3% p<0.01), and excitatory motor neurons (<12% p<0.001), were significantly underrepresented in TS-SOD1 animals (**Figure 2D**). In contrast, transcriptionally active cells (>10% p<0.001), and glial cells (>15%, p<0.001) were proportionally increased in the trisomic mice. This imbalance suggests that Sod1 overexpression may impair the transition of progenitors into mature neurons, either through delayed differentiation or altered lineage fate decisions, while favoring gliogenesis. This pattern is consistent with observations from amyotrophic lateral sclerosis (ALS) models, where SOD1 mutations reduce motor neuron survival both in vivo and in vitro, while glial cells—particularly astrocytes—are relatively spared or even maintained within the central nervous system (Di Giorgio et al., 2007; Meyer et al., 2014). These findings suggest that the neuronal vulnerability and relative glial resilience observed in our SOD1 trisomy model may reflect a shared underlying biology, wherein toxic gain-of-function effects in ALS and gene overexpression in Down syndrome converge to produce similar cell type-specific phenotypes.

We then examined *Sod1/SOD1* transcript levels across the identified ENS cell types. In most populations, TS-SOD1 cells showed the expected ∼1.5-fold increase in total *Sod1* expression relative to WT, reflecting trisomic gene dosage **(Figure 2E**). However, two populations deviated from this trend: interneurons exhibited only a modest 1.2-fold increase (p<0.001), and transcriptionally active progenitors showed a slightly elevated 1.6-fold increase (p<0.001). These differences suggest cell-type-specific regulation of transgene expression or transcript stability and may point to unique vulnerabilities of interneurons to *Sod1* dosage imbalance.

To distinguish transcripts derived from the human transgene from those of the endogenous mouse *Sod1*, we quantified the ratio of human SOD1 to mouse Sod1 expression across ENS cell types (**Figure 2F**). For this purpose, the human SOD1 transcript (ENST00000270142.11) was appended to the mouse reference transcriptome as an additional entry, with its genomic location annotated to the targeted insertion site at the Gt(ROSA)26Sor locus (ENSMUSG00000086429; Chr 6:113,043,843–113,055,336; GRCm39). This annotation allowed us to unambiguously identify, and count reads mapping to the transgene as human (*hSOD1*) and to distinguish them from those mapping to the endogenous mouse gene (*mSod1*). This separation was essential for accurately determining cell-type–specific contributions of transgenic versus endogenous Sod1 expression, given their high sequence identity and the need to resolve potential dosage effects from each source.

Based on the transgene design and expected gene dosage, the anticipated ratio of hSOD1 to mSod1 was approximately 1:2 across all cell types. This pattern was indeed observed in most neuronal populations. Strikingly, however, glial cells deviated from this expectation, exhibiting a significantly higher relative expression of endogenous *Sod1*, with an average *mSod1:hSOD1* ratio of 3.2 (**Figure 2F**). This suggests a glia-specific amplification of mouse Sod1 expression, potentially driven by cell-intrinsic enhancer activity, differential mRNA stability, or post-transcriptional regulation. Such disproportionately high Sod1 expression in glia may contribute to their relative expansion and could exert non–cell-autonomous effects on neighboring neurons, further compounding the ENS defects observed in TS-SOD1 mice.

To corroborate these transcriptomic findings at the protein level, we performed immunofluorescence staining of P0 distal colon sections using the pan-neuronal marker Tubb3 and antibodies against SOD1. Quantification of fluorescence intensity revealed a ∼1.3-fold decrease (p<0.001) in Tubb3 signal in TS-SOD1 mice relative to WT controls (**Figure 2G**), consistent with the observed depletion of neuronal cell types in the single-cell data. SOD1 protein levels were increased by ∼1.2-fold (p<0.001) in the same tissue, confirming transgene overexpression. Notably, cells co-expressing Tubb3 and SOD1 showed a ∼1.25-fold reduction (p<0.001) in intensity in TS-SOD1 mice, suggesting that even neurons expressing SOD1 are reduced in abundance or differentiation status, potentially due to oxidative or metabolic stress induced by Sod1 overexpression.

Together, these data demonstrate that Sod1 trisomy disrupts normal ENS development by reducing the production or maintenance of neuronal subtypes, expanding glial populations, and altering both the magnitude and specificity of Sod1 expression across cell types. The effects are particularly pronounced in inhibitory neurons and glia, offering a cellular framework for understanding how SOD1 contributes to HSCR risk in the context of trisomy 21.

### Sod1 trisomy disrupts gene regulatory networks and suppresses Ret expression in the developing enteric nervous system

To understand the transcriptional consequences of Sod1 trisomy on enteric nervous system (ENS) development, we performed differential gene expression analysis on scRNA-seq data obtained from the distal colon of TS-SOD1 and wild-type (WT) neonates at postnatal day 0 (P0). The ENS cells were computationally extracted and compared between genotypes using a pseudobulk approach (Squair et al., 2021). Differential expression analysis revealed widespread transcriptomic changes in TS-SOD1 mice, with 1878 (FC ≥1.5, FDR≤10^−5^) genes significantly upregulated and 1099 genes significantly (FC ≤1.5, FDR≤10^−5^) downregulated compared to WT (**Figure 3A**).

**Figure 3:**
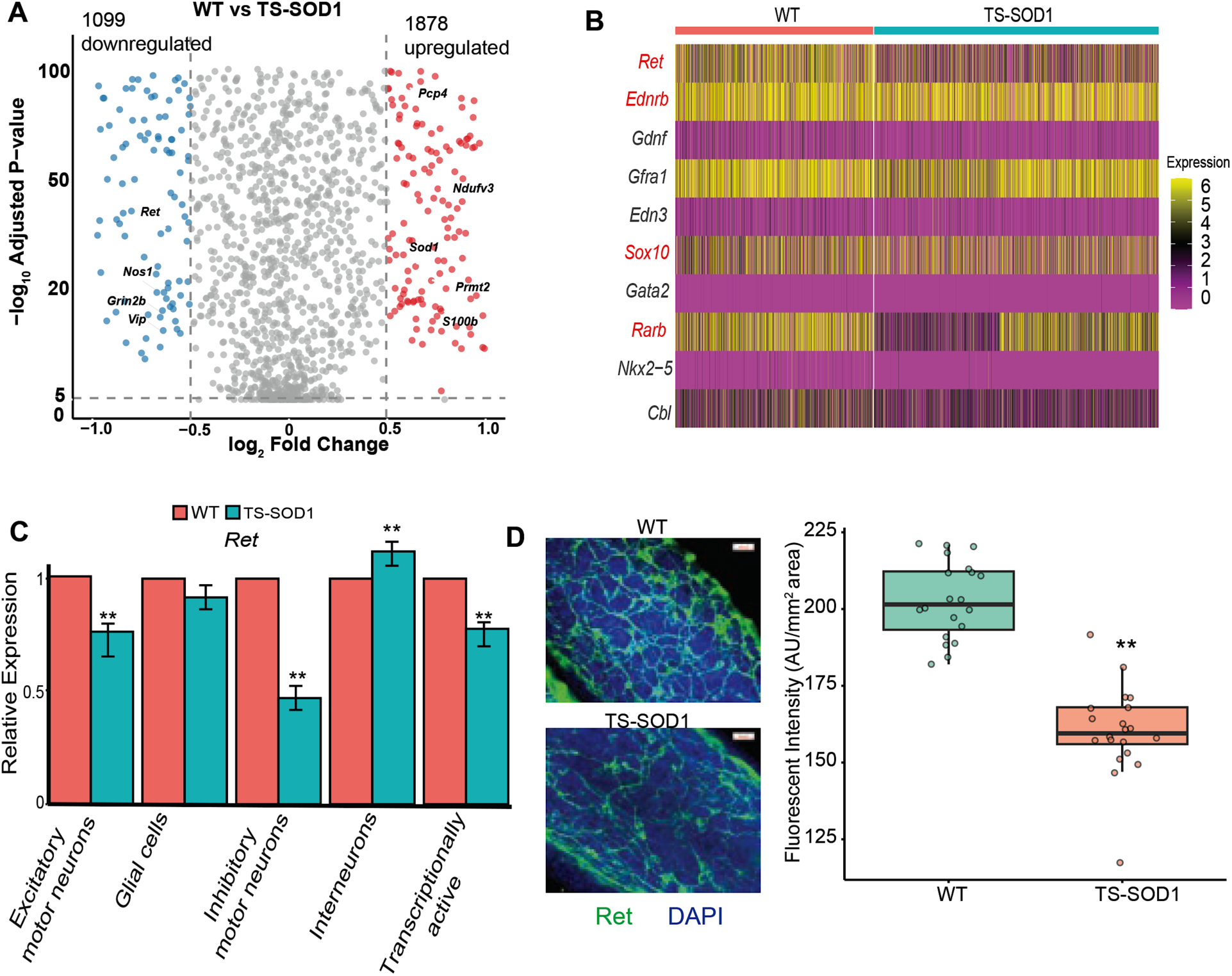
*Sod1/SOD1* trisomy alters expression of HSCR-associated genes in the neonatal ENS. (A) Volcano plot of differential gene expression from scRNA-seq analysis of ENS cells in WT vs. TS-*Sod1/SOD1* mice at P0 with1,878 significantly upregulated and 1,099 downregulated genes in trisomic mice. Upregulated genes include mouse homologs of human Chr21 genes such as *Pcp4, Ndufv3, Prmt2, S100b, and Sod1*, while downregulated genes include the key neuronal markers *Ret, Vip, Nos1, and Grin2b*. (B) Heatmap of the gene expression of 10 genes from a core gene regulatory network associated with Hirschsprung disease. The ligands *Gdnf* and *Edn3* (mesenchymal-specific) and transcription factors *Gata2* and *Nkx2-5* are not detected. Among expressed genes, *Ret, Ednrb, Sox10*, and *Rarb* show reduced expression in trisomic ENS cells. (C) Cell-type–specific analysis of *Ret* expression reveals significant downregulation in excitatory motor neurons, inhibitory motor neurons, and transcriptionally active cells of TS-*Sod1/SOD1* mice. *Ret* expression is mildly but significantly increased in interneurons but unchanged in glial cells. (D) Immunofluorescence quantification of RET protein in distal hindgut shows significantly reduced intensity in TS-*Sod1/SOD1* as compared to WT mice, confirming reduced protein expression *in vivo*. Data are presented as mean ± SEM; statistical significance (p < 0.01) was determined by MAST GLM framework in *Seurat* or using t-tests where appropriate.

Among the downregulated genes, Gene Ontology (GO) enrichment analysis revealed a strong overrepresentation of pathways essential for synaptic function and neuronal connectivity. The most significantly enriched terms included signal release from synapse, neurotransmitter secretion and transport, synapse assembly, and the synaptic vesicle cycle (FDR < 1×10⁻¹³; **Supplementary Figure 1A**). These categories also encompassed processes regulating synapse structure and organization, as well as vesicle-mediated transport within synapses. The coordinated downregulation of these pathways suggests that SOD1 trisomy may impair fundamental mechanisms of neuronal communication in the ENS, potentially contributing to the observed deficits in specific neuronal subtypes. Notably, *Ret, Vip, Nos1,* and *Grin2b* were among the most significantly reduced transcripts. These genes are hallmarks of differentiated enteric neurons, with *Ret* encoding a receptor tyrosine kinase essential for ENS progenitor survival and migration, *Vip* and *Nos1* labeling inhibitory neurons, and *Grin2b* involved in NMDA-type glutamate signaling. The coordinated downregulation of these neuronal markers’ points toward a broader impairment in neuronal maturation and subtype specification in TS-SOD1 mice.

In addition, the downregulated gene set showed enrichment for pathways related to DNA replication and cell cycle regulation, indicating impaired proliferative capacity. This suggests that reduced proliferation may underlie the loss of specific neuronal subtypes, consistent with mechanisms reported in other HSCR models (Fries et al., 2025; Vincent et al., 2023).

In contrast, GO enrichment analysis of the upregulated genes revealed a strong signature of proliferative and genomic maintenance programs, including double-strand break repair via homologous recombination, DNA-templated DNA replication, cell cycle checkpoint signaling, chromosome segregation, and regulation of cell cycle phase transitions (FDR < 1×10⁻¹⁸; **Supplementary Figure 1B**). This enrichment is consistent with a shift toward proliferative progenitor or glial states within the ENS. Notably, Sod1 is known to regulate redox homeostasis and protect dividing cells from oxidative stress (Sturtz et al., 2001; Tsang et al., 2014), thereby supporting cell cycle progression; *Sod1* trisomy may therefore enhance these proliferative programs, contributing to the glial expansion observed in our single-cell data. Among the upregulated genes, five were mouse orthologs of genes found on HSA21 (*Pcp4, Ndufv3, Prmt2, S100b, and Sod1*), consistent with gene dosage effects and successful transgene expression. These include regulators of calcium signaling (Pcp4), mitochondrial function (Ndufv3), and glial maturation (*S100b*), indicating that diverse cellular processes are impacted by the additional *Sod1* allele.

Finally, to specifically evaluate the effect of Sod1 trisomy on known Hirschsprung disease (HSCR)-associated gene networks, we focused on a panel of 10 genes comprising a core RET-EDNRB regulatory axis. This gene set includes ligands (*Gdnf, Edn3*), receptors (Ret, Ednrb), co-receptors (*Gfra1*), transcriptional regulators (*Sox10, Gata2, Nkx2-5, Rarb*), and ubiquitin ligase (*Cbl*). As expected, the ligand genes *Gdnf* and *Edn3*, which are predominantly expressed in mesenchymal or epithelial compartments, were undetectable in the ENS-derived neural crest cells (**Figure 3B**). Similarly, transcription factors *Gata2* and *Nkx2-5* were not expressed at this stage, consistent with their restricted roles in early ENCDC development (Chatterjee and Chakravarti, 2019; Chatterjee et al., 2016). Among the remaining genes, several showed genotype-dependent changes. *Ret, Ednrb, Sox10,* and *Rarb* were significantly (p<0.001) downregulated in TS-SOD1 mice. The suppression of *Ret and Ednrb*—both essential for ENS progenitor proliferation and migration—suggests that *Sod1/SOD1* trisomy may directly interfere with core signaling pathways critical for neuronal survival and maturation reiterating our observation from tissue level expression profiling (**Figure 1C**). *Sox10*, a key transcription factor required for early ENS specification and maintenance of progenitor identity, was also reduced, implying a depletion or dysfunction of neural crest–derived precursors. The downregulation of *Rarb*, a mediator of retinoic acid signaling, further suggests impairment in differentiation pathways that normally promote the transition to mature neuronal fates in ENS.

To explore these effects in a cell-type-specific context, we examined *Ret* expression across defined ENS subtypes (**Figure 3C**). In TS-SOD1 mice, *Ret* was significantly downregulated in excitatory motor neurons (0.5-fold p<0.001), inhibitory motor neurons (1.3-fold p<0.001), and transcriptionally active progenitor-like cells (0.5-fold p<0.001). These are the major neurogenic populations in the ENS at P0 and are particularly sensitive to disruptions in Ret signaling. In contrast, *Ret* expression was mildly (0.3-fold, p<0.001) but significantly increased in interneurons, possibly reflecting a compensatory or context-specific response. Glial cells, which do not depend on Ret signaling for their differentiation, showed no significant change in *Ret* expression, serving as an internal control. This analysis confirms that the reduction in *Ret* expression is not uniform but instead targets neuronal subtypes most vulnerable to HSCR-associated genetic insults.

To determine whether these transcriptional changes translated to protein-level deficits, we performed immunofluorescence staining for RET in the distal colon. Quantification of RET fluorescence intensity revealed a 0.81 (19%; p<0.001 ) reduction in intensity of Ret signal in the distal gut in TS-SOD1 tissues compared to WT controls (**Figure 3D**), validating the transcriptomic findings and confirming a physiologically relevant decrease in Ret protein levels in vivo. Overall, these results provide compelling evidence that Sod1 trisomy disrupts the expression of multiple ENS genes, including critical components of the RET-EDNRB signaling network, and leads to a cell-type–specific reduction in Ret expression. These disruptions likely contribute to the impaired differentiation of enteric neurons and may underlie the increased susceptibility to HSCR observed in trisomy 21.

### Sod1 trisomy alters glial subtype identity and abundance through a Ret-independent mechanism

To further investigate the expansion of the glial compartment observed in TS-SOD1 animals (**Figure 2D**), we performed a focused analysis of glial cells within the ENS at postnatal day 0 (P0). We performed reclustering on the 976 glial cells (401 from wild-type and 575 from TS-SOD1 mice) extracted from our ENS manifold using Seurat’s graph-based approach(Stuart et al., 2019). Clustering was based on 238 high variable genes and the first 15 principal components, which together accounted for 98% of the variance within the glial population. This revealed four transcriptionally distinct glial subtypes (**Figure 4A**), which we annotated as microglia, astrocytes, transcriptionally active glia, and proliferating glia based on unsupervised clustering and marker gene expression profiles (**Figure 4B**). Differential expression analysis identified key genes distinguishing each subtype, including canonical markers such as *S100b, Plp1,* and *Nfia*, as well as subtype-enriched genes like *Hmga2, Camk1d, Mapk10*, and Prdx1.

**Figure 4:**
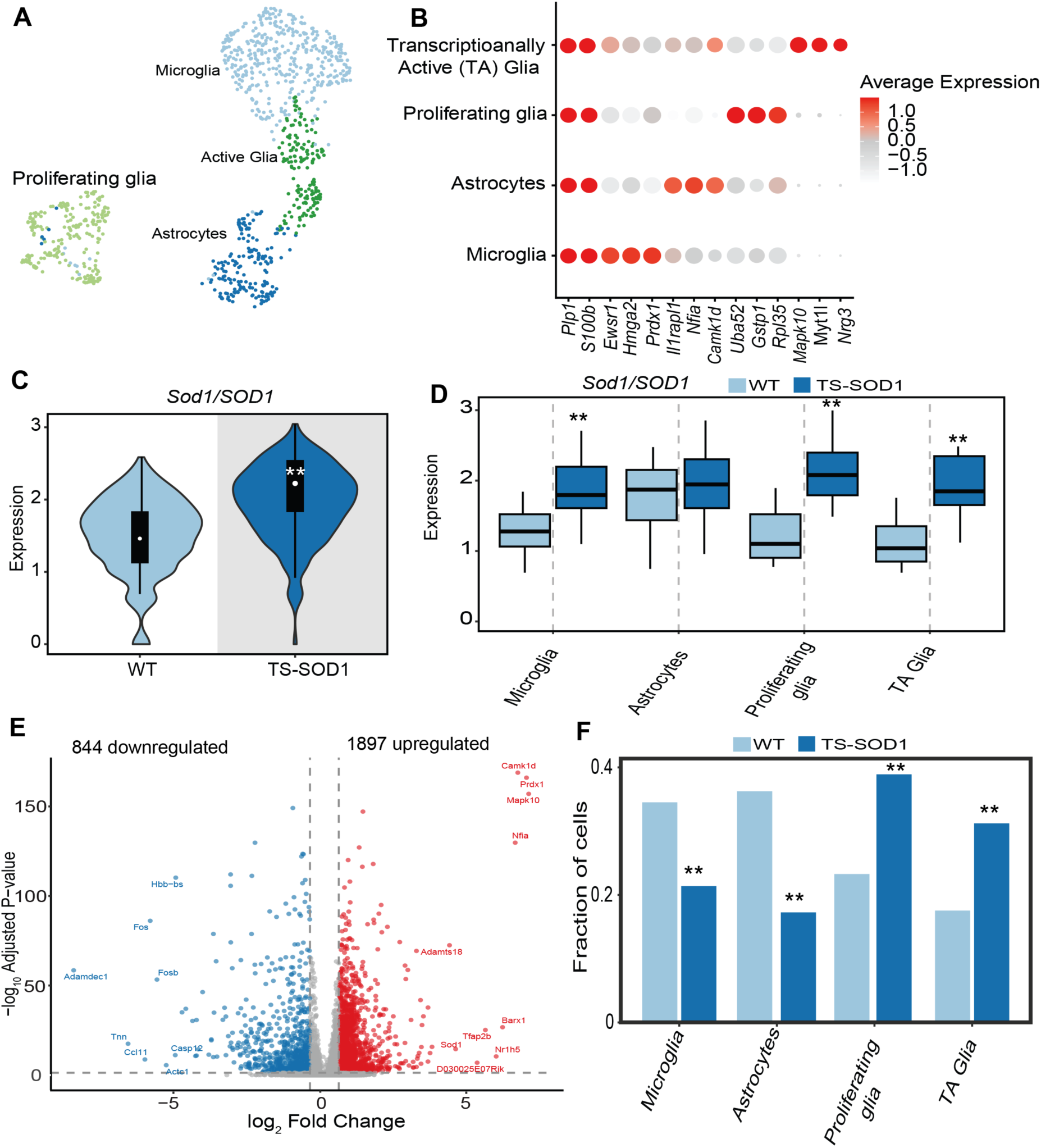
Sod1 trisomy alters glial subtype composition and transcriptional identity in the neonatal ENS. (A) Reclustering the glial cell population from scRNA-seq identifies its four distinct subtypes (microglia, astrocytes, active glia, and proliferating glia) based on differential gene expression. (B) Heatmap of the top marker genes for each glial subtype, including *Plp1, S100b, Hmga2, Prdx1, Il1rapl1, Nfia, Camk1d, Mapk10*, etc., confirms subtype identity. (C) Average *Sod1/SOD1* expression is elevated ∼1.6-fold in glial cells of TS-*Sod1/SOD1* relative to WT mice. (D) Subtype-specific analysis showing ∼2-fold higher *Sod1/SOD1* expression in proliferating and active glia, ∼1.5-fold in microglia, but no significant change in astrocytes. (E) Volcano plot showing differentially expressed genes in glial cells between genotypes. TS-*Sod1/SOD1* glia shows 1,897 significantly upregulated and 844 downregulated genes. (F) Bar plot of glial subtype proportions reveals increased abundance of proliferating and active glia in the TS-*Sod1/SOD1* ENS, and reduced proportions of astrocytes and microglia as compared to WT ENS. Data are presented as mean ± SEM; statistical significance (p < 0.01) was determined by MAST GLM framework in *Seura*t or two-way ANOVA.

We next assessed the expression of *Sod1/SOD1* within the glial compartment. On average, glial cells from TS-SOD1 mice expressed Sod1 at 1.5-fold higher levels than WT, consistent with the trisomic gene dosage (**Figure 4C**). However, this elevation was not uniform across subtypes (Figure 4D). Proliferating glia and transcriptionally active glia showed the highest overexpression (∼2-fold), while microglia showed a moderate ∼1.5-fold increase. Notably, astrocytes exhibited no change in *Sod1* expression compared to WT, suggesting that the impact of Sod1 trisomy on glial gene dosage is highly context-dependent and may selectively influence immature or reactive glial states.

To evaluate the functional consequences of Sod1 overexpression on glial transcriptional identity, we conducted differential expression analysis comparing TS-SOD1 and WT glia. This revealed widespread transcriptional remodeling, with 1897 genes significantly upregulated and 844 genes downregulated in the trisomic condition (**Figure 4E**).

Gene Ontology (GO) enrichment analysis of the upregulated genes in glial cells revealed strong activation of proliferative and genomic maintenance pathways, including DNA-templated DNA replication, double-strand break repair via homologous recombination, recombinational repair, cell cycle checkpoint signaling, and mitotic cell cycle phase transitions (FDR < 1×10⁻⁹; **Supplementary Figure 2A**). These terms indicate an overall bias toward cell cycle re-entry and genomic stability mechanisms within the glial population. Among the upregulated genes were *Camk1d, Mapk10, Prdx1, and Nfia*, all of which are implicated in glial activation, stress responses, or proliferation. The enrichment of nuclear transport and nucleocytoplasmic transport pathways further suggests an upregulation of biosynthetic and signaling processes required to sustain active cell growth and division, consistent with the observed expansion of proliferating and active glia in TS-SOD1 mice.

Gene Ontology (GO) enrichment analysis of the downregulated genes in glial cells highlighted processes associated with extracellular matrix organization, extracellular structure organization, cell–substrate adhesion and connective tissue development (FDR < 3×10⁻¹³; **Supplementary Figure 2B**). Many of these pathways are critical for maintaining structural integrity, cell–cell communication, and the supportive functions of glia within the ENS. Conversely, several genes typically associated with glial homeostasis or neuronal interaction—such as *Fos, Tnn, and Casp12*—were downregulated, indicating a shift away from a quiescent or supportive phenotype. This expression profile is consistent with a more activated, proliferative, and potentially gliotic transcriptional state, as reflected in the concurrent upregulation of cell cycle and DNA replication pathways in the same cell population and highlights a balance required by the glial cells to proliferate without.

We then assessed the proportional representation of each glial subtype across genotypes. In TS-SOD1 mice, there was a significant expansion of proliferating and transcriptionally active glia, whereas astrocytes and microglia were proportionally reduced (**Figure 4F**). These findings suggest that Sod1 trisomy promotes a shift in glial fate toward immature or reactive states, potentially at the expense of differentiated, homeostatic glia. This remodeling of glial subtypes parallels the reduction of neuronal lineages observed earlier and suggests a broader imbalance in ENS lineage allocation or maturation. Importantly, since *Ret* expression is affected mature enteric glia in TS-SOD1 glial populations (**Figure 3C**), this indicates that the observed glial expansion and transcriptional remodeling occur independently of Ret signaling, and therefore likely represent a distinct cellular consequence of Sod1 overexpression.

Taken together, these findings reveal that Sod1 trisomy alters the composition and molecular identity of glial cells in the ENS through a Ret-independent mechanism. The expansion of proliferative and active glia, accompanied by elevated Sod1 expression and global transcriptomic changes, suggests an additional non-neuronal route by which trisomy 21 may contribute to ENS dysfunction and disrupted neuronal-glial homeostasis.

## Discussion

Down Syndrome, like many other developmental disorders, is associated with many congenital disorders affecting numerous organ symptoms. Although the panoply of phenotypes associated with this trisomy must involve many genes the genetic origin of its specific sub-phenotypes is murky. There is no doubt that trisomy for HSA21 is the primary risk factor but which other genes are necessary to lead to the specificity of each sub-phenotype. Since not all DS individuals have every sub-phenotype where does the sequence basis for this difference lie? In this study we answer this question for Hirschsprung disease (HSCR) which occurs at a 100-fold increased frequency in DS individuals (Amiel et al., 2008; Badner et al., 1990; Torfs and Christianson, 1998).

Here we demonstrate that the HSCR association in DS arises from increased dosage of *SOD1* interacting with *RET*, one of two major genes for non-syndromic HSCR. The studies reported here are based on a targeted humanized mouse transgene comprising the human *SOD1* gene (hSOD1) inserted in the mouse *Rosa26* locus yielding a trisomy from 2 copies of the endogenous mouse *Sod1* gene (mSod1) and one copy of the human transgene. This mouse model demonstrates that *SOD1* may be sufficient to induce defects in enteric neurogenesis that are intrinsic to HSCR. However, despite the cellular defects observed in the mouse distal colon at birth (P0), in humans, the aganglionosis in HSCR requires hypomorphic mutations in *RET* as well (Arnold et al., 2009; de Pontual et al., 2007). These findings taken together confirms our hypothesis that increased dosage of *SOD1* on HSA21 interacts with *RET* to further reduce its gene expression in individuals who already harbor the hypomorphic *RET* enhancer variant leading to <50% *RET* gene expression and aganglionosis (Fries et al., 2025). Thus, syndromic HSCR and non-syndromic HSCR may all arise from a central underlying defect in RET signaling.

By isolating SOD1 dosage in an otherwise euploid genome and resolving consequences at single-cell resolution in the developing gut, this study demonstrates that one HSA21 gene can recapitulate key features of the DS–HSCR axis. Importantly, the approach does more than nominate SOD1 as “associated”: it establishes sufficiency for perturbing the enteric nervous system (ENS) through two separable routes—*Ret*-dependent neuronal defects and *Ret*-independent glial remodeling. Conceptually, this provides a template for deconstructing DS phenotypes one candidate at a time, asking (i) whether increased dosage is sufficient to perturb a given developmental program and (ii) which cell types, pathways, and time windows are most vulnerable.

The SOD1 model reveals that T21-level dosage does not act uniformly across cell types. In neurons, excess SOD1 converges on attenuation of Ret signaling—a central axis for enteric neurogenesis—providing a mechanism for reduced production or maintenance of specific neuronal subtypes implicated in HSCR. In glia, by contrast, SOD1 dosage promotes proliferative and reactive states without altering *Ret,* yielding a *Ret*-independent expansion of particular glial subpopulations. These orthogonal outcomes—*Ret* suppression in neurons and proliferative bias in glia—highlight how a single dosage change can have cell-type-specific effects driven by distinct regulatory networks and stress responses.

The divergence between neuronal loss and glial expansion resonates with broader neurobiology: neurons, with high metabolic demand and limited redox buffering, are preferentially injured by oxidative imbalance, whereas glia more readily activate stress-adapted, proliferative programs. The parallels between amyotrophic lateral sclerosis (ALS) and our SOD1 trisomy model support a convergent pathophysiological framework: whether via toxic gain-of-function mutations in ALS or overexpression due to trisomy 21, SOD1 dysregulation generates a redox-imbalanced environment that preferentially injures neurons while allowing glia to survive or expand. In the ENS, this imbalance manifests as *Ret*-dependent suppression of neurogenic lineages and *Ret*-independent proliferation of reactive glial subtypes. This dual mechanism could help explain why individuals with Down syndrome are more susceptible to HSCR and other ENS-related pathologies—neuronal deficits arise directly from impaired RET signaling, while altered glial composition may exacerbate or perpetuate network dysfunction through non–cell-autonomous effect.

The single-gene sufficiency shown here argues that DS-associated congenital disorders can be parsed into “dosage-sensitive nodes” (e.g., SOD1) embedded within canonical developmental pathways (RET/EDNRB, guidance, cell-cycle checkpoints). This has several implications. First, it motivates systematic testing of additional HSA21 candidates—alone and in rational combinations—to map which nodes are sufficient for which syndromic traits. Second, it prioritizes modifier loci and environments that interact with those nodes (e.g., RET coding/enhancer variation, semaphorin signaling, oxidative stress buffering), providing hypotheses for variable penetrance in DS. Third, it suggests potential therapeutic angles that are pathway-specific rather than chromosome-wide—for example, strategies that bolster RET signaling in vulnerable neuronal lineages or temper reactive gliosis without broadly suppressing glial support functions.

Technically, combining a humanized allele at a permissive locus with single-cell multi-compartment readouts (transcript dosage partitioning, trajectory inference, and protein validation) offers a general blueprint for DS biology. The ability to distinguish transgene-derived versus endogenous transcripts and to quantify dosage at cell-type resolution was critical for uncovering the glia-specific amplification of mouse Sod1 and for separating RET-dependent from RET-independent effects. Deployed across other DS-enriched conditions—congenital heart defects, immune dysregulation, craniofacial anomalies—this design can turn diffuse “trisomy effects” into testable causal circuits.

As with any targeted model, ectopic insertion may not fully capture native regulatory context, and neonatal analyses provide a developmental snapshot rather than a temporal continuum. Future experiments that (i) titrate SOD1 dosage, (ii) combine SOD1 trisomy with sensitizing Ret enhancer or coding variants, (iii) test cell-type-restricted Sod1 manipulations, and (iv) probe oxidative and proteostatic stress pathways in vivo will refine causality and therapeutic leverage points. Parallel work should evaluate whether normalizing SOD1 dosage rescues neuronal deficits without exacerbating glial dysfunction, and whether modulating RET/RA or semaphorin signaling can buffer SOD1-driven risk in the ENS.

DS is a syndromic condition in which organ-specific vulnerabilities emerge from selective dosage-sensitive nodes interacting with core developmental pathways and background modifiers. By demonstrating that SOD1 alone can recreate hallmark ENS disturbances along RET-dependent and RET-independent axes, this study provides both mechanistic insight into the DS–HSCR connection and a broadly applicable strategy for dissecting the genetic architecture of DS-associated syndromes—one candidate, one tissue, and one cell state at a time.

## Materials & Methods

### Generation of SOD1 trisomic mice

Generation of humanized SOD1 mice was achieved by pronuclear microinjection of strain JR#036152 (C57BL/6J-Gt(ROSA)26Sor^em7Mvw^/MvwJ), which contained dual heterologous Bxb1 attachment sites in the ROSA26 loci (Low et al., 2022)to allow for Bxb1 mediated recombinase-mediated cassette exchange (RMCE). The microinjection mix comprised of 100 ng/µl BXB1 mRNA (Low et al., 2022) and 3.5 ng/µl of modified Ch17-349N04 BAC donor dna in 10 mM Tris w/ 0.1mM EDTA). The BAC Ch17-349N04 (PAC biosciences) was modified using two rounds of Red/ET Recombination (Genebridges) to insert Bxb1 attB sites. The first round of recombineering used a PCR amplicon containing Ampicillin and the Bxb1 attB-GT sequence (GGCTTGTCGACGACGGCGGTCTCCGTCGTCAGGATCAT) with 33 bp long homology arms at each end to place the insert 470 bp upstream of SOD1-DT (ENST00000449339.1) in the BAC. The second round of recombineering used a PCR amplicon containing aminoglycoside phosphotransferase from Tn5 and the Bxb1 attB-GT sequence (GGCTTGTCGACGACGGCGGACTCCGTCGTCAGGATCAT) with 33 bp long homology arms at each end to place the attB site 8037 bp downstream of SOD1. The BAC was isolated using the QIAGEN Large-Construct Kit following manufacturer’s instructions, further purified using a phenol:chloroform:isoamyl alcohol extraction and was then subjected to drop dialysis using 25 mm diameter, Type-VS Millipore membrane (Millipore, Inc) floating on 10 mM Tris w/ 0.1mM EDTA for two hours.

### Genotyping

To verify correct insertion of the human SOD1 locus into the mouse ROSA26 site, we used a three-primer PCR strategy. The primers included a ROSA26 forward primer (5′ AAT GCC AAT GCT CTG TCT AGG 3′) and a ROSA26 reverse primer (5′ GTC GCT CTG AGT TGT TAT CAG T 3′), which amplify the endogenous ROSA26 allele, together with a human SOD1–specific reverse primer (5′ CAC AGA ATT GGG CAA CTA TCA C 3′), which yields a product only when the hSOD1 sequence is present at the locus. In mice carrying a single-copy insertion of hSOD1, PCR amplification produced two bands, an ∼800 bp fragment corresponding to the targeted hSOD1 allele and an ∼700 bp fragment corresponding to the wildtype ROSA26 allele. In contrast, wildtype mice showed only the ∼700 bp ROSA26 band, confirming correct and specific insertion of the human SOD1 construct.

### Phenotype

TS-SOD1 mice were born at the expected Mendelian ratios but, by 8 weeks of age, weighed 7.36 ± 0.41 g (mean ± SD, n = 4) compared to 12.30 ± 0.30 g (mean ± SD, n = 4) in wild-type littermates. Despite the reduced body weight, TS-SOD1 mice exhibited normal feeding behavior and general activity. The lower weight may reflect impaired digestive efficiency secondary to ENS disruption.

### Dissection and dissociation of embryonic tissue

Foregut and hindgut tissue from Δmcs+9.7/Δmcs+9.7; +/+ and Δmcs+9.7/Δmcs+9.7; CFP/+ at E12.5 and E14.5 were dissociated into single-cell suspensions using Accumax (Sigma, USA). The cells were filtered serially through a 100 μm and 40 μm cell strainer, and centrifuged at 2,000 rpm for 5 m. The cell pellets were resuspended in 5% FBS, 4 mM EDTA in Leibovitz L-15 medium. This cell suspension was diluted to 20,000 cells and processed through the 10X Genomics GEM generator to create a standard 3’ library as per the 10X Genomics standard protocol. These cells were sequenced at an average of 1×10^5^ ± 10^4^ reads/cell.

### Single cell gene expression (RNA-seq) analysis

Sequence files were processed through *CellRanger* to generate the *barcodes, features* and *matrix* files. These files were processed through *Seurat* (Stuart et al., 2019)to generate a matrix count file. Individual datasets for each genotype were then analyzed by normalizing and scaling the data followed by detecting high variable genes across cell types for downstream analysis using *sctransform* in *Seurat* (Hafemeister and Satija, 2019). Cells which met the following criteria were retained for further downstream analysis: (nUMI >= 500) & (nGene >= 250) & (log10GenesPerUMI > 0.80) and (mitoRatio < 0.20)). We only retained those genes which were detected in >10 cells. To eliminate technical differences between datasets and to perform comparative scRNA-seq analysis across experimental conditions we utilized *Integration* analysis in *Seurat* which matched shared cell populations across datasets. These methods first identify cross-dataset pairs of cells that are in a matched biological state (‘anchors’), before creating clusters of these cells to avoid bias. These are used as input to the uniform manifold approximation and projection (UMAP) dimensionality reduction tool to establish a biologically relevant two-dimensional embedding of the cells. Analysis pipeline used for *Seurat* can be found at https://github.com/SumantraChatt/mouse-scRNAseq/tree/main

### Single cell differential gene expression analysis

We used Seurat’s *FindMarker* feature to perform differential expression by pseudo-bulking the samples either across genotypes or across cell clusters. We utilized “MAST”: GLM-framework (Finak et al., 2015) that treats cellular detection rate as a covariate within *Seurat*.

### Cell-type annotation

We annotated cell types by performing differential gene expression analysis by considering all markers with >1.5-fold change (FDR<0.05) in that cluster, as compared to all other clusters. These genes were analyzed for their GO terms using DAVID (Huang et al., 2007) and enriched GO terms (FDR <0.01 and containing >10 genes) were used to annotate the clusters. Additionally we used *clusterprofiler* (v 4.16.0) (Yu et al., 2012) to annotate genes into biological processes from both GO and Kyoto Encyclopedia of Genes and Genomes (KEGG) pathways for conformation of the primary annotation using DAVID. For enteric nervous system cluster, along with using unbiased cell type marker identification we also used previously identified markers for specific neuronal and glial cell types in the ENS (Elmentaite et al., 2021; Morarach et al., 2021; Vincent et al., 2023).

### Immunofluorescence assays on whole mouse gut

Distal colon tissue at P0 were dissected and washed in ice-cold PBS followed by fixation in 4% PFA at 4°C for 1 h. The fixed tissues were permeabilized and blocked in 5% normal donkey serum and 0.3% Triton-X for 1 h at room temperature (RT). The tissues were incubated with appropriate primary antibodies at 4°C overnight, washed with PBS, then incubated in appropriate secondary antibodies at 4°C overnight. Tissues were washed in PBS and incubated with 0.2 μg/mL of DAPI (4′,6-diamidino-2-phenylindole) for 10 m at RT, followed by incubation for 20 m at RT with CUBIC-1 (16) diluted 1:10 with PBS. Samples were washed with PBS and mounted on Fisherbrand Superfrost Plus microscope slides with ProLong Gold Antifade mounting media and imaged using a Zeiss AxioVert Microscope.

### Image Analysis

The following workflow was used in ImageJ to analyze fluorescent images, using previously described methods for neuronal density (Berk-Rauch et al., 2023). Fluorescence intensity of specific markers (TUBB3, SOD1, RET) was used as a proxy for neuronal density within each hindgut region. For each image, the area positive for marker staining was quantified and expressed as a percentage of the total area of the region of interest. Genotype information was blinded to the observer during analysis. Data are presented as mean marker intensity across the region of interest from multiple biological replicates per genotype. For double fluorescence analysis (TUBB3 and SOD1), the intensity of each fluorescent marker was quantified in ImageJ. Regions where both markers co-localized within the same cell were identified and used for further analysis. Data are presented as the average intensity of each marker within co-expressing regions across the entire distal colon, averaged over multiple embryos per genotype. Statistical significance was assessed using a paired Student’s *t-*test.

### Gene expression Taqman assays

Total RNA was extracted from distal colon at P0 cells using TRIzol (Life Technologies, USA) and cleaned on RNeasy columns (QIAGEN, USA). 500μg of total RNA was converted to cDNA using SuperScript IV reverse transcriptase (Life Technologies, USA) using Oligo-dT primers. The diluted (1/5) total cDNA was subjected to Taqman gene expression (Thermo Fisher Scientific) using transcript-specific probes and primers for *Sod1* (Mm01344233_g1; Thermo Fisher Scientific). Mouse β-actin was used as an internal loading control, as appropriate, for normalization. Three independent mice were used for RNA extraction and each assay performed in triplicate (n = 9).

The mean of the ΔC_t_ values (C_t_ Condition - C_t_ Actin) for each replicate for each condition was calculated after converting them to a linear scale (2^ΔCt^). The fold change was calculated as the ratio of the mean of 2^ΔCt^ experiment / 2^ΔCt^control. The mean for the control was set to unity and the relative fold change was calculated based on these values. For calculating the variance of the of the fold change we used the following formula: 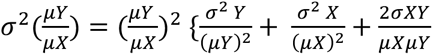

where *σ*^2^ is variance, *μ* is mean and 2*σXY* is the covariance between an experimental condition Y and its corresponding control X. Subsequently, P-values were calculated from pairwise 2-tailed t tests, and the data presented as the fold change with its standard error.

## Author Contributions

SC and AC conceived and designed the study. JR and AZ generated the TS-SOD1 mice. GG and LW conducted single cell and immunofluorescence assays; SC analyzed the data. SC and AC wrote the manuscript. All authors approved the final version of the manuscript.

