## Supplementary Figure 1 and 2 for "*Sod1* trisomy causes ENS developmental defects and susceptibility to Hirschsprung disease via neuronal *Ret* suppression and glial remodeling"

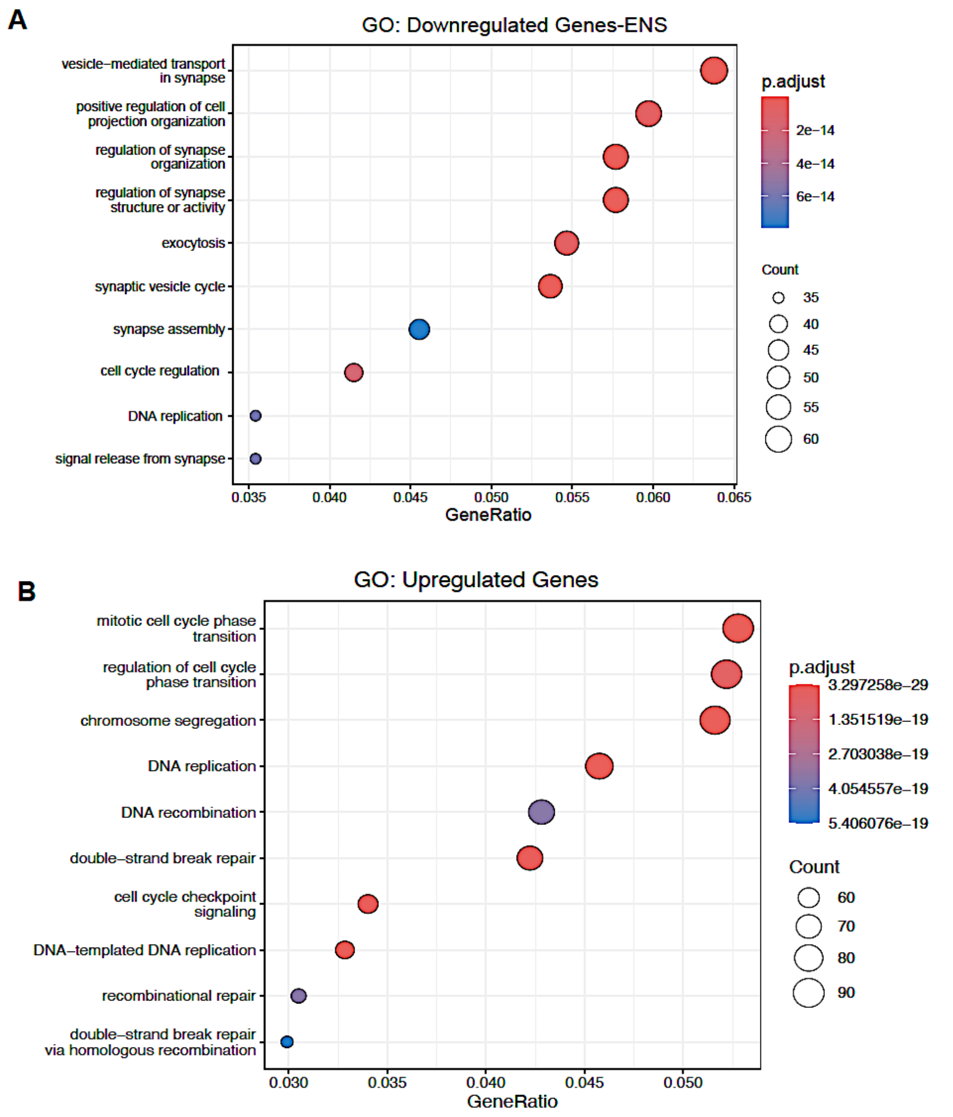


**Supplementary Figure 1: GO annotation of upregulated and downregulated genes in the ENS of TS-SOD1 mice.**

(A) The 1,878 downregulated genes in the ENS of TS-SOD1 P0 mice cluster into 10 significantly enriched biological processes, dominated by synaptic and neuronal functions including synaptic vesicle cycle, exocytosis, vesicle mediated transport in synapse, synapse assembly, regulation of synapse organization or activity, and positive regulation of cell projection organization, indicating impaired neuronal connectivity and neurotransmission.

(B) Upregulated genes are enriched for cell cycle and DNA maintenance pathways, including DNA replication, chromosome segregation, mitotic cell cycle phase transition, cell cycle checkpoint signaling, and double strand break repair, consistent with aberrant activation of proliferative and genome integrity programs.

Bubble size reflects gene count, color denotes adjusted p value, and the x axis indicates gene ratio

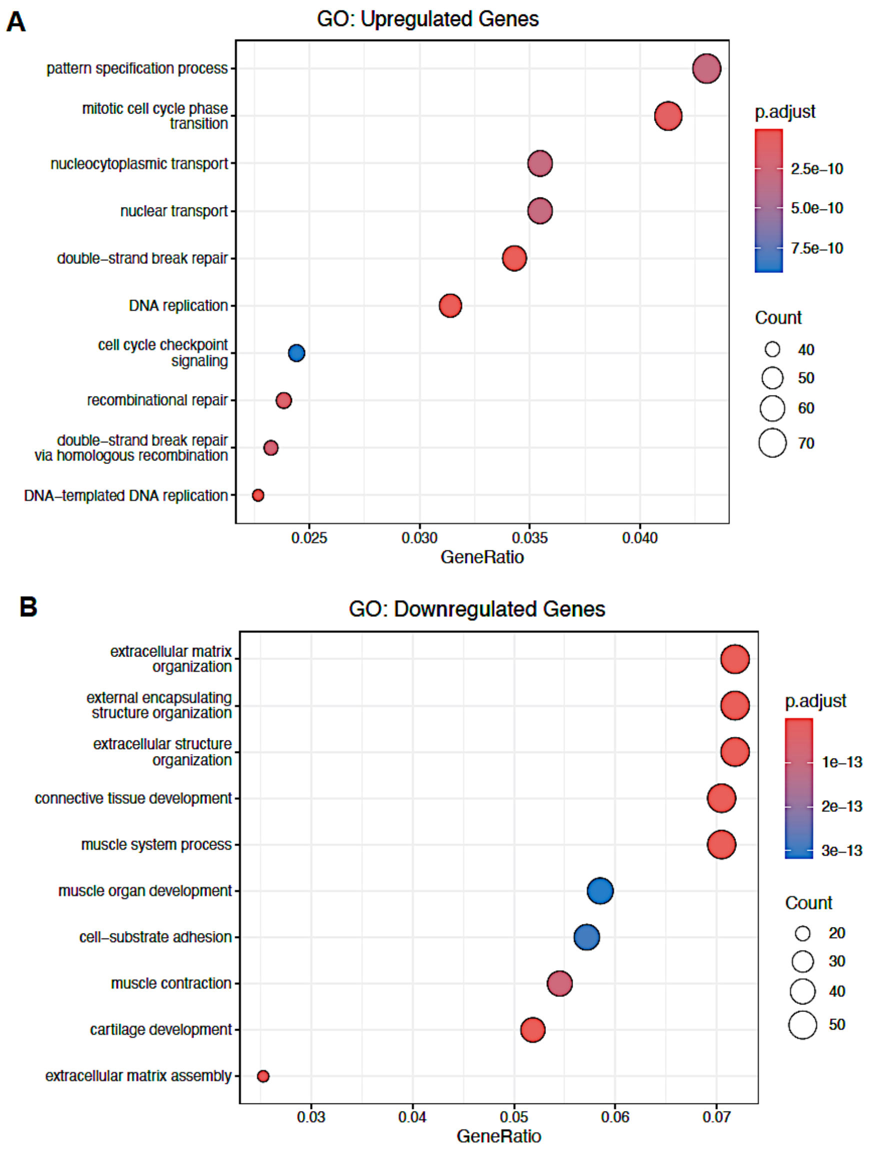


**Supplementary Figure 2: GO annotation of differentially expressed genes in enteric glial cells of TS-SOD1 mice compared to wildtype.**

(A) Genes upregulated in the glial population of TS-SOD1 P0 mice are enriched for cell cycle and genome maintenance pathways, including DNA replication, mitotic cell cycle phase transition, cell cycle checkpoint signaling, nuclear and nucleocytoplasmic transport, and double strand break repair, indicating aberrant activation of proliferative and DNA damage response programs in glia.

(B) Downregulated glial genes are significantly associated with extracellular matrix organization, extracellular structure and external encapsulating structure organization, cell substrate adhesion, and connective tissue and muscle related developmental processes, consistent with impaired glial interactions with the extracellular niche.

Bubble size denotes gene count, color represents adjusted p value, and the x axis indicates gene ratio
